# Optimizing Rhamnolipid Biosynthesis: Evaluating Predictive Methods using *Pseudomonas aeruginosa* Mutants

**DOI:** 10.64898/2026.01.15.699715

**Authors:** Ingrid Yoshimura, Jonas Contiero, Eric Déziel

## Abstract

Rhamnolipids (RLs) are versatile biosurfactants produced by *Pseudomonas aeruginosa* with significant industrial potential. However, high production costs remain a barrier to large-scale use, necessitating genetic strategies to improve yields. While many genes are reported to influence RL production, studies often rely on qualitative phenotypic assays of questionable reliability. We systematically evaluated 29 *P. aeruginosa* PA14 mutants using traditional assays (Siegmund-Wagner blue plates, swarming motility) and validated the findings using Liquid Chromatography-Mass Spectrometry (LC/MS). We found that traditional phenotypic assays have a high misprediction rate (∼35-38%), primarily due to confounding factors like variable flagellar function, colony spreading, and growth rates. Specifically, LC/MS quantification revealed that *rpoN* and *pvdQ* knockouts significantly increased total rhamnolipid titers, whereas *crc*, *dksA*, and *dspI* knockouts decreased production. Notably, the increased titers in *rpoN* and *pvdQ* mutants were linked to enhanced biomass accumulation rather than higher per-cell biosynthetic rates. These findings highlight the critical necessity of using quantitative analytical methods for accurate strain screening and provide a clarified set of genetic targets for metabolic engineering aimed at optimizing rhamnolipid production.

**Importance:** This study addresses a critical methodological flaw in biosurfactant research: the over-reliance on qualitative phenotypic assays that too often lead to inaccurate conclusions. By systematically comparing traditional screening methods with LC/MS quantification across a collection of *Pseudomonas aeruginosa* mutants, we demonstrate that common assays like swarming motility and blue plates fail to accurately predict rhamnolipid production in over one-third of cases. These inaccuracies lead to the misidentification of genetic targets and may waste resources in metabolic engineering efforts. Our work provides a reliable framework for strain screening, identifies specific genes that influence rhamnolipid yields, and clarifies the biological factors—such as flagellar motility and growth dynamics—that bias traditional results. These findings are essential to optimize biosurfactant production and ensure data reproducibility.

## Introduction

Biosurfactants are microbially derived compounds increasingly explored as sustainable alternatives to conventional chemical surfactants (1). They are synthesized by microorganisms such as bacteria and offer several advantages, including high biodegradability, low toxicity, production from renewable resources, structural diversity, selectivity, and stability under extreme conditions of temperature and salinity (2–4). Among them, rhamnolipids (RLs) stand out as glycolipid biosurfactants with remarkable properties and demonstrated commercial potential. However, large-scale applications would still benefit from increased production yields (5, 6). This challenge is driving efforts to develop high-producing microbial strains. *Pseudomonas aeruginosa*, the main natural RL producer, regulates RL biosynthesis, especially the *rhlAB* biosynthetic genes, through a complex regulatory network involving three interconnected quorum sensing (QS) systems (7). Due to this high level of regulatory complexity, many genes could influence RL production directly or indirectly. For instance, many factors are involved in social swarming motility and biofilm formation - processes closely tied to RL production (8–13) - suggesting that previously poorly investigated genetic alterations may impact RL production.

To explore this relationship, a literature review was conducted to list genes that have been identified to influence RL production and/or regulatory pathways, either positively or negatively. The results of this survey are summarized in **Table 1**. Although numerous studies have reported that knocking-out specific genes leads to altered RL production, the methods commonly used to assess such changes are varied and their reliability to reach such conclusion may be questionable (14). For instance, a frequent approach is the use of swarming motility assays on semi-solid agar plates as a proxy for RL production by *P. aeruginosa*. However, any correlation between swarming behavior and RL levels should be expected inconsistent as this phenotype is also dependant on flagellar function. In the present study, we used 29 mutants from a nonredundant library to evaluate the reliability of various methodologies and identify their limitations in RL production assessments.

**Table 1.**
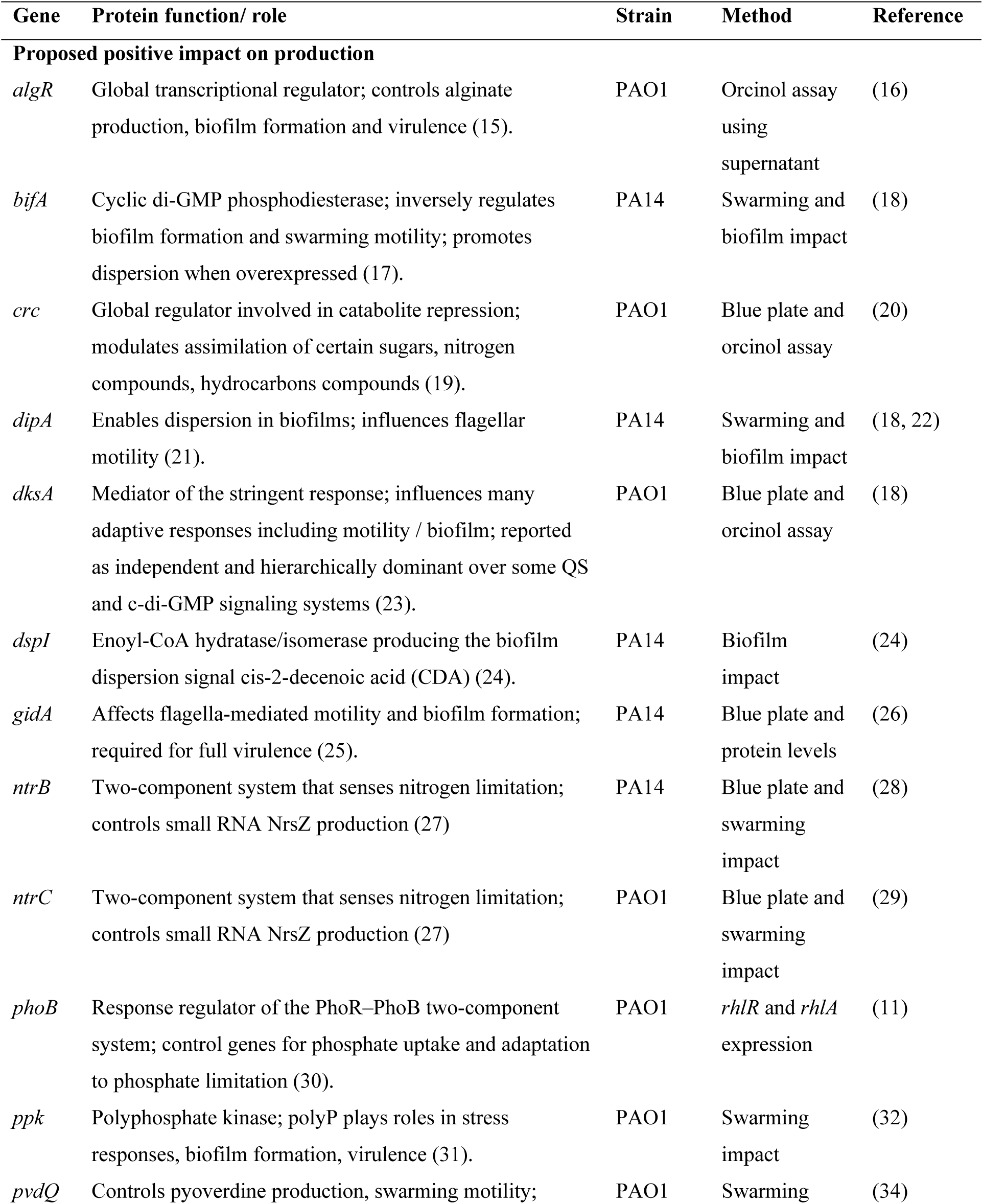
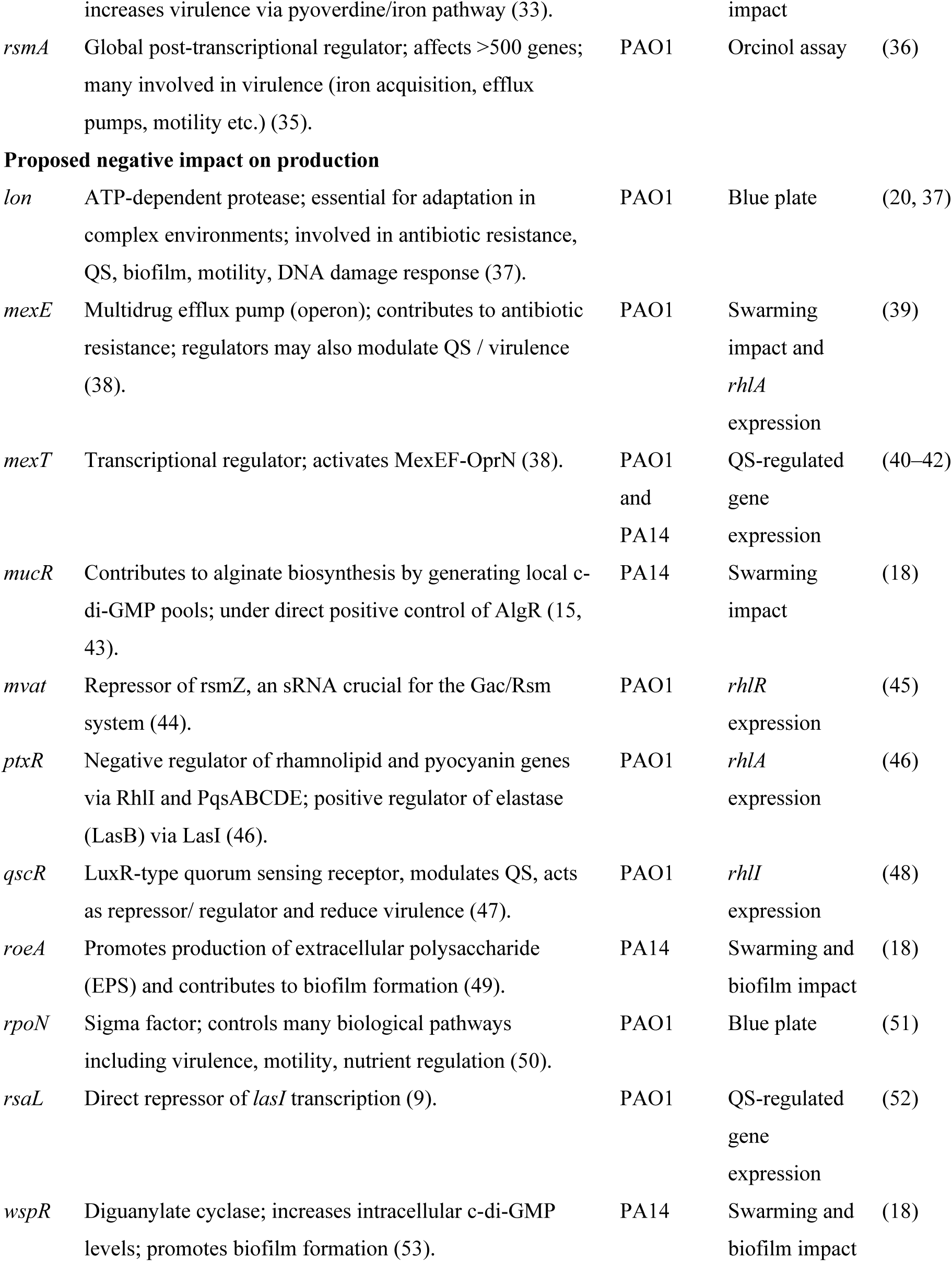
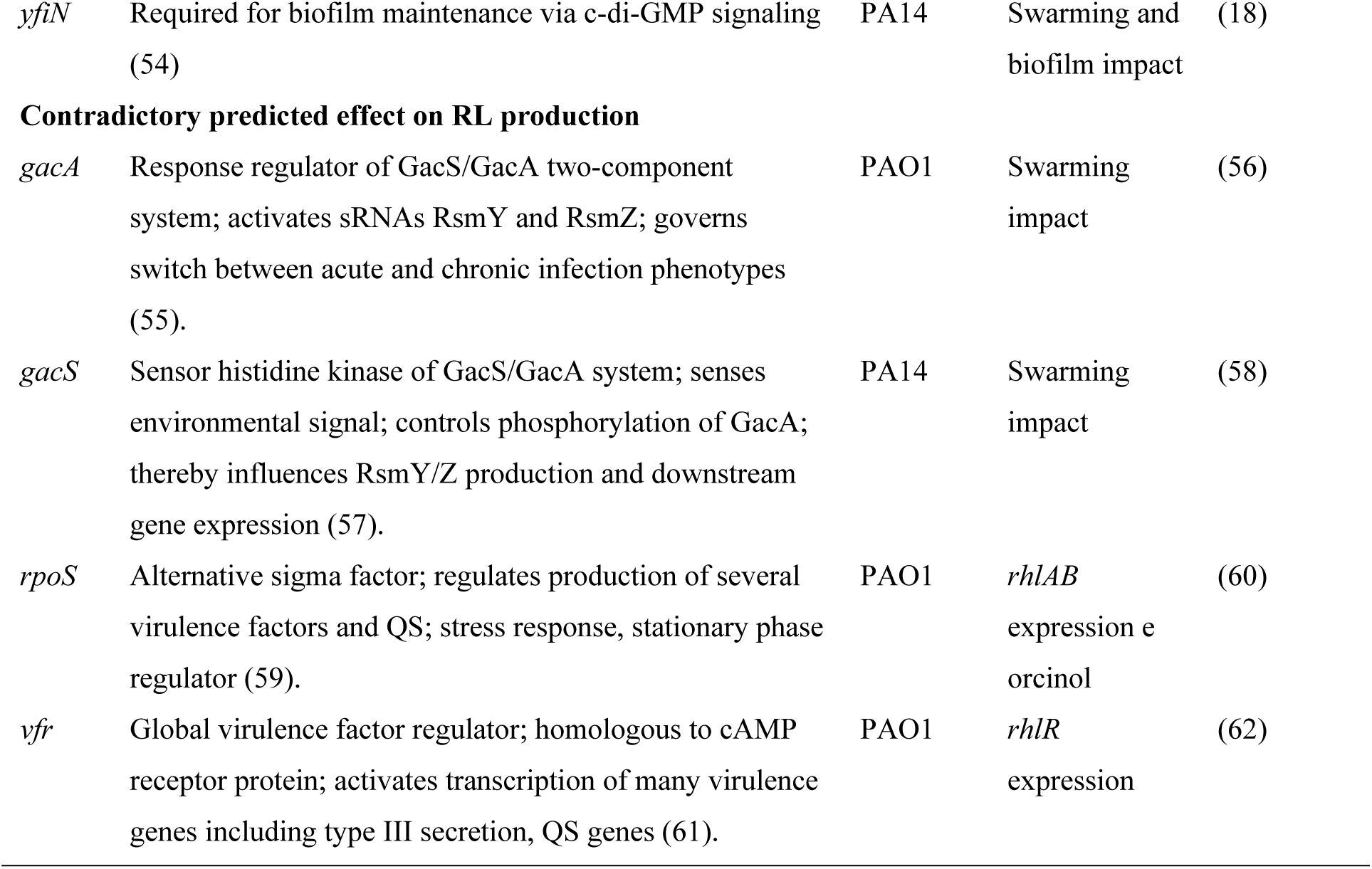
Genes reported to influence rhamnolipid production in *Pseudomonas aeruginosa*.

## Results and Discussion

### Assessing swarming motility as a predictive method for rhamnolipid production

Over the years, several genes have been implicated in the production of RLs and/or the expression of the *rhlAB* biosynthetic genes across different *P. aeruginosa* strains, under various culture conditions and using diverse methods. In order to clarify the literature and reconcile often-contradictory findings, we compared several knockout mutants of the prototype strain PA14 side-by-side (listed in Table 1), under the same experimental conditions and using the same methodologies.

As an initial indicator of differences in biosurfactant production, a swarming assay was performed. Swarming motility is a rapid and coordinated multicellular movement across a semi-solid surface, driven by morphological differentiation and intercellular interactions (63). A distinctive feature of *P. aeruginosa* swarming is the formation of dendritic, fractal-like tendrils radiating from the inoculation site (64), a process associated with the production of RLs and its precursor 3-(3-hydroxyalkanoyloxy) alkanoic acids (HAAs) (65). It is therefore commonly used as an indirect measure of differential RL synthesis. As shown in **Figure 1**, the *lon* and *crc* mutants exhibited no swarming under our experimental conditions. Mutants for *bifA*, *dipA*, *dksA, gacA, gacS, pvdQ, rpoN* and *rsmA* showed severely reduced swarming motility, while *rsaL* and *dspI* displayed mild swarming compared to wild-type (WT) PA14. Mutants in genes such as *algR*, *gidA*, *mexE*, *mexT*, *mucR*, *ntrC*, *phoB*, *ppk*, *ptxR*, *qscR*, *roeA*, *vfr*, *wspR*, and *yfiN* exhibited swarming patterns similar to WT. None produced obviously enhanced swarming motility, suggesting no mutant would produce enhanced levels of RL compared to the parental background.

**Figure 1.**
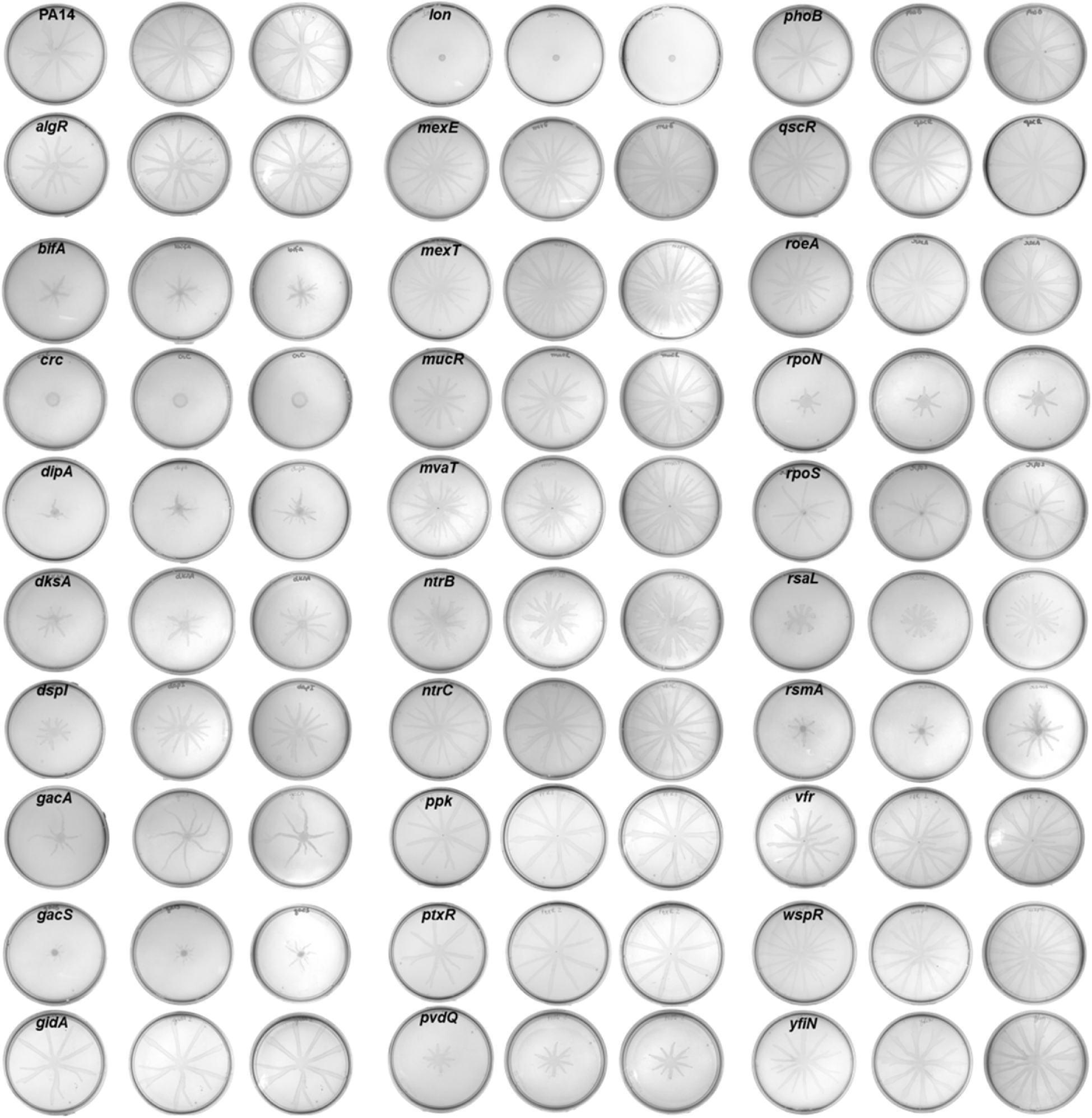
Swarming motility assays of *P. aeruginosa* PA14 and various isogenic mutants. Strains were assessed after 16-20 h of incubation. Assays were performed in triplicates with the wild type included for comparison to validate the reproducibility of the results.

Based on these results and following the hypothesis that swarming motility correlates with RL production, it would be predicted that *crc* and *lon* mutants are defective in biosynthesis of this wetting agent. In *P. aeruginosa*, deletion of *crc* disrupts the *rhl* quorum sensing system, which would thereby impair RL synthesis since it positively regulates *rhlAB*. Crc, along with the RNA chaperone Hfq, normally represses the expression of the Lon protease, which targets RhlI, the enzyme responsible for producing the Rhl quorum sensing signal *N*-butanoyl-L-homoserine lactone (C_4_-HSL). In the absence of Crc, Lon levels increase, leading to RhlI degradation, reduced C_4_-HSL synthesis, and consequently, downregulation of RL production (20). Notably, deletion of *lon* or constitutive expression of *rhlI* restores RL levels, confirming the role of this regulatory cascade (20). Thus, the *crc* knockout is expected to reduce or inhibit RL production, whereas the *lon* knockout should not. This raises the question: is the lack of swarming motility of the *lon* mutant really explained by the absence of RL production? Additionally, swarming motility requires a functional flagellum (64, 65).

Therefore, in addition to RL production, we need to consider whether altered flagellar regulation might explain some of the swarming phenotypes observed. Indeed, several selected mutants are involved in the regulation of the second messenger bis-(3′-5′)-cyclic dimeric guanosine monophosphate (c-di-GMP), a key regulatory molecule in bacteria typically associated with biofilm formation and a sessile lifestyle (66, 67). In general, low levels of c-di-GMP are linked to motility and virulence factor production, whereas high levels suppress flagellar activity and promote the expression of adhesins and biofilm-associated exopolysaccharides (EPSs) (18, 66). Accordingly, it is reasonable to assume that mutants with a tendency to form more biofilm may exhibit reduced flagellar activity and, consequently, diminished swarming motility. The *dipA* mutant (dispersion-induced phosphodiesterase A) exhibits increased biofilm formation and reduced swarming motility (68), findings that align with our observations under the tested conditions. In *P. aeruginosa* PA68, inactivation of *dipA* was associated with increased biofilm formation, likely due to compromised flagellar function (18). Specifically, DipA loss significantly inhibits flagellar motor switching and directional reversal during swimming in a MapZ-dependent manner (21). In this case, less swarming would not be a consequence of reduced RLs.

Taken together, differences in swarming behavior compared to WT can be attributed to one or more of the following factors: [a] altered growth rate (e.g. some mutants may have growth defects or poor adaptation to the swarming medium), [b] modified RL production, and/or [c] different motility (e.g. flagellar defects could result in reduced spreading). Next, we verified these factors separately.

### Comparative growth analysis of mutant strains

Considering that growth should also be considered when assessing the RLs production potential of a bacterium, we next assessed whether any of the mutants exhibited growth defects in the M9DCAA culture medium and monitored their growth for 36 h with a Bioscreen system. This medium is the same used for swarming assays, without agar. The results revealed that the *rsaL* and *lon* mutants display impaired growth under these conditions (**Figure S1**). The inability of the *lon* mutant to swarm may thus be explained by its growth defect, while the *rsaL* mutant exhibited reduced swarming compared to the wild type, which could also be partially attributed to its compromised growth. In contrast, the *gacS*, *dipA*, *bifA*, *rsmA*, and *rpoS* mutants demonstrated enhanced growth compared to the WT under the same conditions. Interestingly, all mutants that exhibited increased growth showed a reduced swarming motility phenotype, as well. Thus, since growth alone did not account for all cases of increased or decreased swarming, we turned to a more specific colorimetric assay on agar plates to better understand whether the observed differences in swarming patterns were related to RL production or not.

### Siegmund-Wagner blue plate assay for rhamnolipid detection

The Siegmund-Wagner blue plates method allows the direct observation of RL production around a colony, by the formation of halos resulting from the interaction of the anionic biosurfactant with a cationic surfactant, causing its precipitation, thus enabling semi-quantitative assessment of RL production (69). This is a widely used method for screening and considered a straightforward and reliable way to verify RL production. However, although apparently simple, we noted that interpretation of results can be challenging because the area of the RL halo zone is affected by the size of the colony; indeed, sometimes, it is not possible to conclude whether the zone is absent or hidden behind the colony **(Fig. 2A)**, and bacteria forming smaller colonies might, misleadingly or not, produce smaller halos. To partially account for these factors, we measured the total diameter of the colony plus the RL production zone, as shown in Figure 2B.

**Figure 2.**
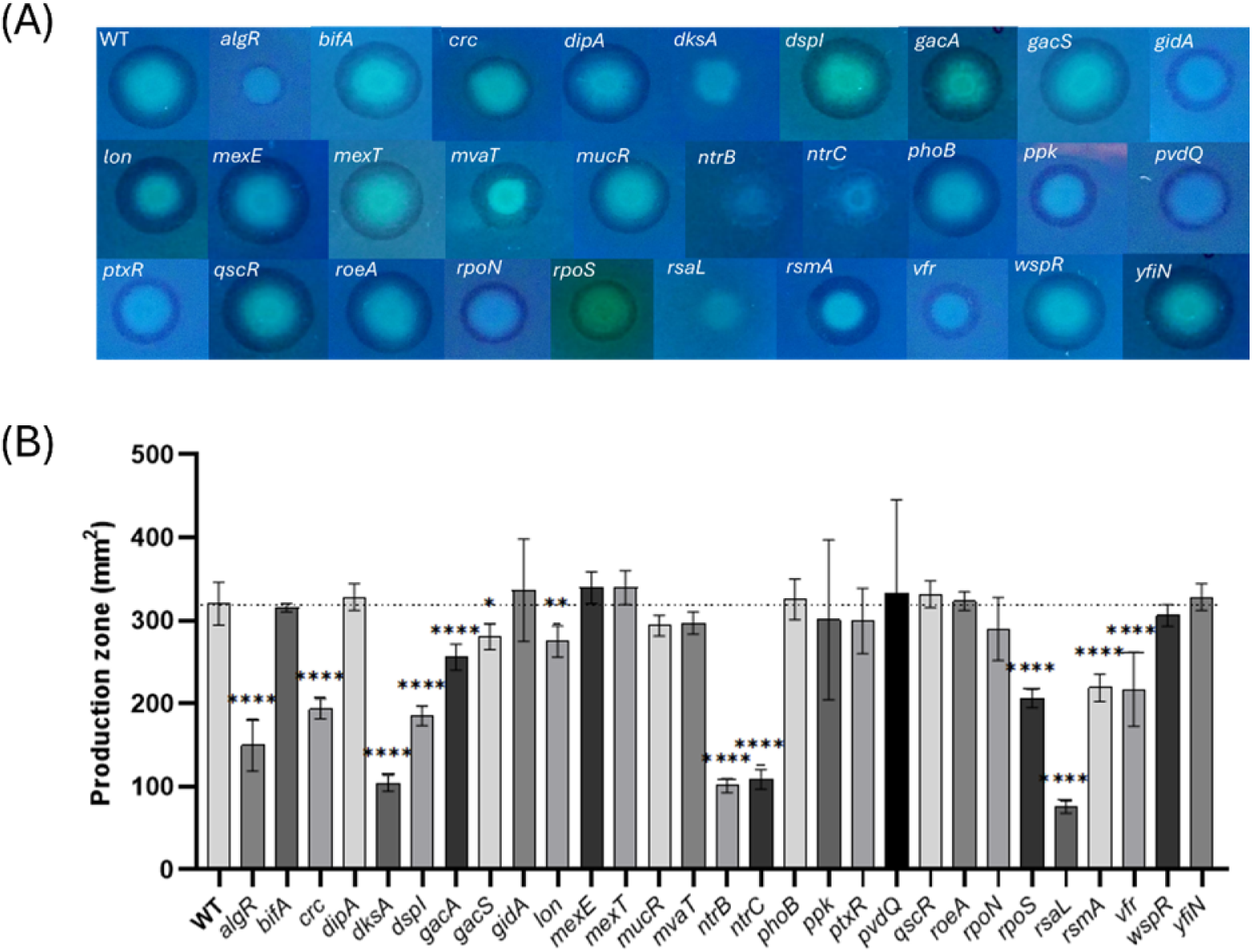
Rhamnolipid production zones by *Pseudomonas aeruginosa* PA14 wildtype and mutants on Siegmund-Wagner blue plates. (A) Pictures of plates after 72h of incubation. (B) Rhamnolipid production zones represented by the mean area of the production zone including the colony. Data were analysed by one-way ANOVA followed by Dunnett’s post hoc test vs. control (\*\*\*\**p* < 0.0001, \*\**p* < 0.01, \**p* < 0.05).

While some mutants, including *dipA*, *gidA*, *mexE*, *phoB*, *pvdQ*, *roeA*, *wspR* and *yfiN* presented halo areas comparable to the WT, none displayed a significantly larger zone, thus presumably higher RL production. Among these mutants indistinguishable from WT, all except *dipA* and *pvdQ* demonstrated swarming behavior also similar to the WT, suggesting that these genes are actually not involved in RL production, at least under the tested conditions. Other mutants such as *dipA* and *pvdQ* displayed defective swarming but WT RL production on blue plates. This adds to the possibility that other factors, such as a flagellar function defect, could play a role — which likely explains the behavior of the *dipA* mutant, as previously discussed.

The mutants *algR*, *crc*, *gacA*, *lon*, *mvaT, ntrB, ntrC, rsaL, rsmA, and vfr* formed smaller colonies on blue plates, which also interferes with interpretation of the results; potential growth defects in this specific condition and/or impaired motility function could be affecting colony size. Among these, only *lon* exhibited impaired growth in the Bioscreen assay, raising the possibility that the reduced colony size of the other mutants might be due to defective flagellar function rather than growth limitation.

The *dksA*, *ntrC*, *ntrB*, and *rsaL* mutants did not form visible halos at all, indicating no RL production under the tested conditions. However, those mutants swarmed in the previous test (Fig. 1), where *dksA* and *rsaL* swarmed less than the WT, but *ntrB* and *ntrC* exhibited swarming patterns similar to WT. This observation suggests two possible hypotheses: RL production is influenced by the composition of the culture medium, which is different in blue plates; or a halo is formed, but its size is too small to be discernible, potentially remaining obscured beneath the bacterial colony.

Colony size influenced halo interpretation, which could again lead to biased conclusions, in the same way as swarming motility: some colonies might spread more easily due to increased flagellar function or better adaptation to the growth medium, not necessarily due to higher RL production. To better understand the impact of flagellar function of mutants on swarming motility and blue plates results, a swimming assay was conducted to investigate potential flagellar activity defects.

### Swimming assay-based evaluation of flagellar motility is needed for interpretation of swarming and blue plates results

Since a swarming defect or smaller colonies can result from defective flagellar motility and thus cause incorrect interpretation of results obtained on plates in the context of RL production, a swimming assay was conducted to investigate potential flagellar function defects **(Figure S2) (Fig. 3)**.

**Figure 3.**
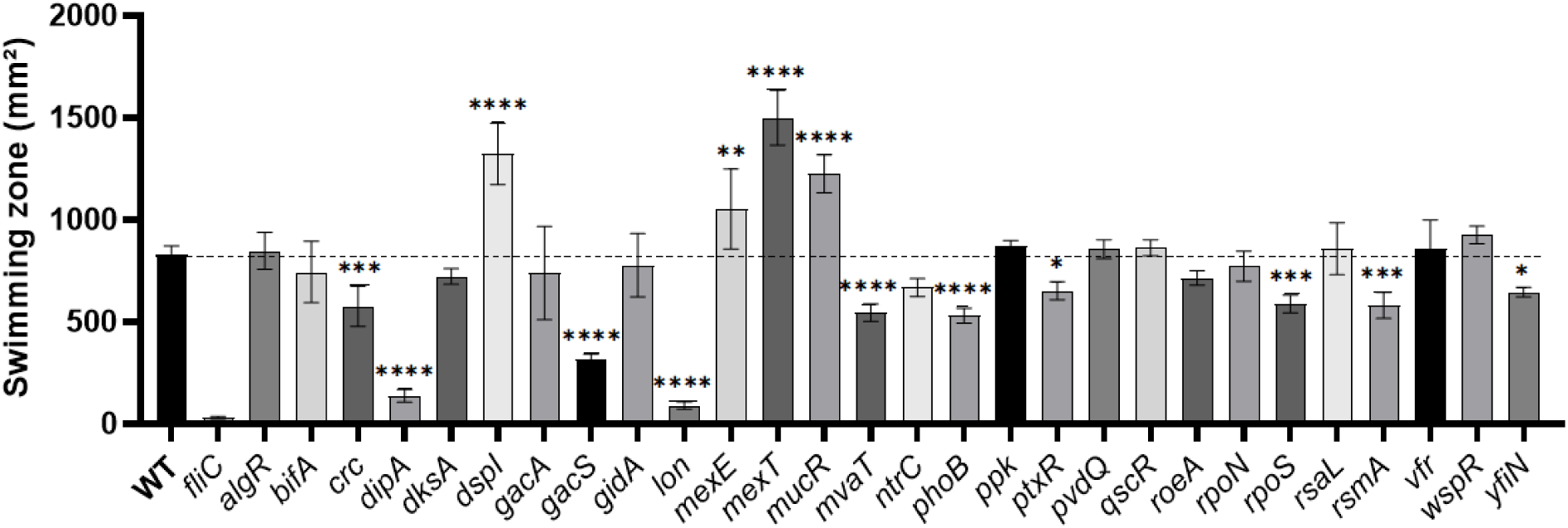
Swimming motility of *P. aeruginosa* PA14 and various mutants. The assay was performed on M9DCAA with 0.25% agar plates. Strains were incubated at 37°C and the measurements were performed after 16 hours. Δ*fliC* mutant was included in the assay as a negative control. Data were analysed by one-way ANOVA followed by Dunnett’s post hoc test vs. control (\*\*\*\**p* < 0.0001, *** *p* < 0.001, \*\**p* < 0.01, \**p* < 0.05).

The *crc*, *dipA*, *gacS*, *lon*, *mvaT*, *ntrC*, *phoB*, *ptxR*, *rpoS*, *rsmA and yfiN* mutants exhibited reduced swimming compared to the WT, while *dspI*, *mexE*, *mexT* and *mucR* showed enhanced swimming. Based on these results, it can be inferred that the reduced swarming motility observed for *crc*, *dipA*, *gacS*, *rpoS*, *lon*, and *rsmA* mutants may be attributed to flagellar defects. Among these, *crc* also displayed reduced RL production, and *lon* showed impaired growth, further contributing to their limited swarming colony size.

On the other hand, *mvaT* and *phoB* demonstrated swarming similar to the WT despite reduced swimming, raising the possibility that these mutants may compensate for defective flagella by increasing RL production. Following this reasoning, mutants *mexE*, *mucR* and *mexT* that exhibited enhanced swarming might have benefited from other mechanisms, such as increased motility, and may in fact produce lower amounts of RL.

However, since flagellar function also influences colony spreading on blue agar plates, and both swarming and halo assays can be biased by growth or motility differences, direct quantification of RL is necessary to validate these hypotheses. The most accurate approach for this is liquid chromatography coupled with mass spectrometry (LC-MS) for direct measurements of molecules in liquid culture samples.

### Measurement of rhamnolipid production by LC-MS quantification

To verify if the swarming and blue plates assay combined with a swimming assay to evaluate flagellar activity allowed for identification of altered RL-producing mutants, we performed LC-MS quantifications. LC-MS analyses are a reliable method to quantify RL levels since it allows for direct quantifications from cultures. For RL production measurements, bacteria were grown in M9DCAA. All mutants were included except *rsaL* and *lon* since they exhibited growth defects **(Fig. S1)**. After five days of cultivation, biomass dry weight measurements indicated that *gacA*, *rpoN*, *dspI*, *wspR*, and *bifA* exhibited higher growth than the WT **(Figure 4A)**. Among these, only *bifA* displayed enhanced growth in the Bioscreen assay after 36 hrs **(Fig. S1)**. The others possibly benefited from a late-phase growth advantage, as the prior assay was limited to a shorter monitoring window, or contributed to biomass beyond cells *per se*, such as EPS or polyhydroxyalkanoate (PHA) granules.

**Figure 4.**
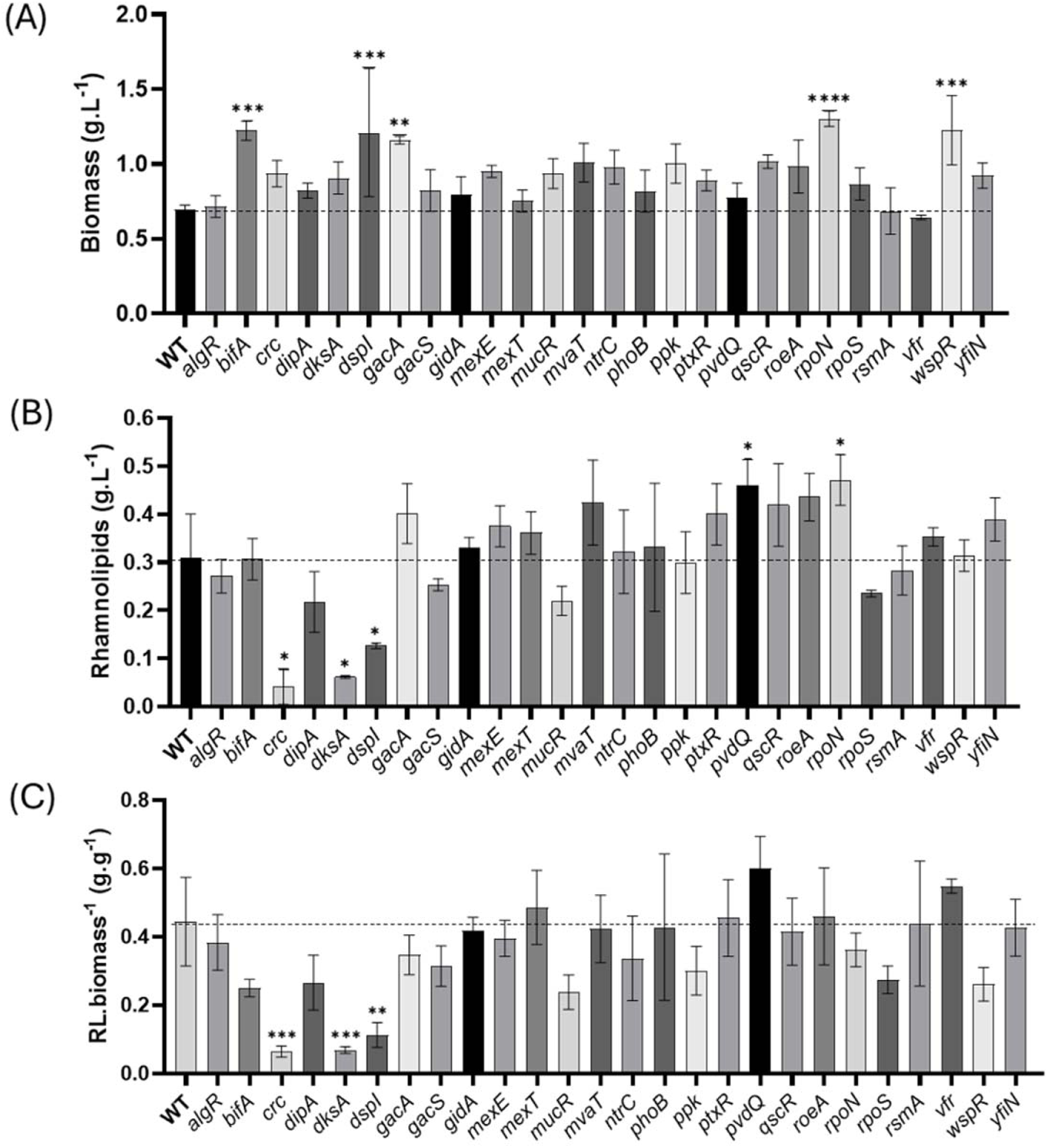
Biomass and rhamnolipid production in *P. aeruginosa* PA14 and mutants. (A) Biomass quantification by dry mass. (B) Rhamnolipids quantification by LC-MS. (C) The strains were cultivated in tubes with liquid M9DCAA supplemented with dextrose for 5 days. Data were analysed by one-way ANOVA followed by Dunnett’s post hoc test vs. control (\*\*\*\**p* < 0.0001, *** *p* < 0.001, \*\**p* < 0.01, \**p* < 0.05).

Regarding total RL production, LC-MS quantification revealed increased production in *pvdQ* and *rpoN* cultures, and reduced levels by the *crc*, *dksA*, and *dspI* mutants **(Fig. 4B).** Although other mutants showed potential for higher RL production, the differences were not statistically significant. However, when production was normalized to cell biomass, the specific yield (RL/biomass) for *rpoN* and *pvdQ* remained comparable to that of the WT **(Fig. 4C)**. This suggests that the increased RL accumulation observed in these mutants results primarily from greater cell growth or a prolonged productive phase, rather than an intrinsic increase in per-cell synthesis rates. From a process perspective, this profile can be advantageous, as higher titers facilitate and reduce the cost of downstream recovery and purification. Nevertheless, from a metabolic-engineering standpoint, *rpoN* and *pvdQ* do not show a real increase in biosynthetic capacity per unit of biomass.

The RpoN sigma factor is a global transcriptional regulator that enables *P. aeruginosa* to rapidly adjust gene expression in response to environmental changes. It governs several processes related to virulence and QS (50). Although RpoN primarily modulates the transcription of metabolic genes, it also regulates key components of the QS network, including *lasI*, *rhlI*, and *pqsR* (51, 70). Our results are in agreement with the hypothesis that RpoN functions as a repressor of the *rhl* QS system by binding to the promoters of *rhlI*, *rhlA*, and *rhlR*. Furthermore, RpoN downregulates the expression of *gacA*, encoding for another major global regulator of virulence factors (36, 51). Nevertheless, other studies have reported that RpoN can act as an activator of QS (60, 70), suggesting that its regulatory role may be context-dependent, which is consistent with previous findings that culture conditions strongly influence the expression of QS-regulated genes (71).

On the other side, the protein PvdQ is involved in the maturation of pyoverdine, a siderophore important for iron acquisition in *P. aeruginosa* (72, 73). Moreover, PvdQ plays a significant role in QS regulation by hydrolyzing acyl-homoserine lactones (AHLs), particularly *N*-3-oxo-dodecanoyl-L-homoserine lactone (3-oxo-C_12_-HSL), thereby influencing the expression of AHL-dependent genes such as *rhlAB* (33, 74). A study explored the use of PvdQ to reduce the virulence of *P. aeruginosa* and demonstrated that treatment with this enzyme inhibited the activity of a *rhlA* promoter-luciferase fusion, suggesting that PvdQ-mediated degradation of QS signals reduces *rhlAB* expression (75). *pvdQ* mutants also displayed increased biofilm formation and reduced swarming motility likely due to signal accumulation (34, 76), and in our study, we found a similar increase in RL production under these conditions. Another work found decreased swarming but no difference in RL production (33).

These findings further highlight that the regulation of swarming motility is not directly linked to RL production, but rather to the broader coordination of multicellular behavior mediated by several factors, including QS. The importance of signal concentration in swarming regulation is a possibility from observations that *pvdQ* mutants, that should be accumulating higher levels of 3-oxo-C_12_-HSL signal, display impaired swarming motility **(Figure 1)** (37). This finding aligns with previous work, which demonstrated that QS signals are required for swarming: *las* system mutations reduce swarming, while *rhl* mutations abolish it entirely (64). These observations suggest that an optimal balance of QS signals is critical, and deficiency or excess of these molecules can both disrupt multicellular coordination. Taken together, PvdQ’s role in swarming motility is not merely a consequence of altered RL levels but likely involves fine-tuned control of QS signal turnover, which is essential for proper spatial and temporal coordination of collective motility.

In a different manner, the *dksA* mutant produced decreased levels of RL and showed less swarming compared to the WT. Jude *et al.* suggested that DksA inhibits RL synthesis by repressing C_4_-HSL production (77). Nonetheless, DksA appears to support basal-level translation of *rhlAB* during the stationary phase. Transcriptional fusion assays showed that *lasB* and *rhlAB* are transcribed normally in the *dksA* mutant, but elastase and RL levels were markedly reduced. This indicates that DksA acts post-transcriptionally, possibly by influencing mRNA translation or protein stability (77).

Clearly, the correlation between swarming behavior and RL levels is not always consistent. Despite the established role of RLs in facilitating swarming, discrepancies are observed between swarming ability and actual RL production levels measured in liquid cultures, which could lead to questioning the accuracy of the swarming assay as a screening method for factors influencing RL production. For instance, for the *gacS* mutant, impaired flagellar function reduces both swimming and swarming motility. Nonetheless, RL production in cultures remains comparable to that of the wild type, as confirmed by LC-MS, indicating that reduced swarming is not linked to an altered RL biosynthesis ability in this case. Similarly, *bifA* mutants exhibit diminished swarming motility, likely due to impaired flagellar reversal mechanisms (17). A reduced frequency of flagellar reversals limits the cell’s ability to change direction efficiently, which is essential for the flexible and coordinated movements required during complex swarming behavior (17). However, swimming remains unaffected, and RL levels are unchanged.

Interestingly, Liu *et al.* demonstrated that reduction in c-di-GMP - specifically through activation of the phosphodiesterase NbdA - triggers a shift from biofilm formation to increased RL production (78). Therefore, decreasing c-di-GMP may act as a molecular switch that favors biosurfactant synthesis, suggesting a promising strategy to enhance RL production by modulating intracellular c-di-GMP levels. In this context, the knockout of genes such as *yfiN*, *roeA*, *mucR*, and *wspR* would be associated with lower c-di-GMP levels and thus expected to enhance RL production; indeed, some of these even produce large swarming zones. Conversely, knocking-out *bifA* and *dipA* were expected to maintain higher c-di-GMP levels and favor biofilm formation (18), thereby presumably reducing RL synthesis. However, in all these mutants, RL production remained similar to the WT strain.

### Limitations and integration of phenotypic prediction methods

In this study we wanted to verify the reliability of classical methods often used in the literature to screen for factors leading to modified RL production or strains with different RL production ability. Based on our results, summarized in **Table S1,** we were able to reach some conclusions on whether swarming motility and Siegmund–Wagner blue plate assays may serve as reliable predictors of RL production in *P. aeruginosa* mutants **(Figure 5)**. This comparison not only assessed the predictive power of these phenotypic methods against known mutant profiles but also provided insight into the biological factors that contribute to inconsistencies and misinterpretations in RL production data in the literature. To this end, both assays were examined for their accuracy and limitations in reflecting the actual biosynthetic capacity of different mutants.

**Figure 5.**
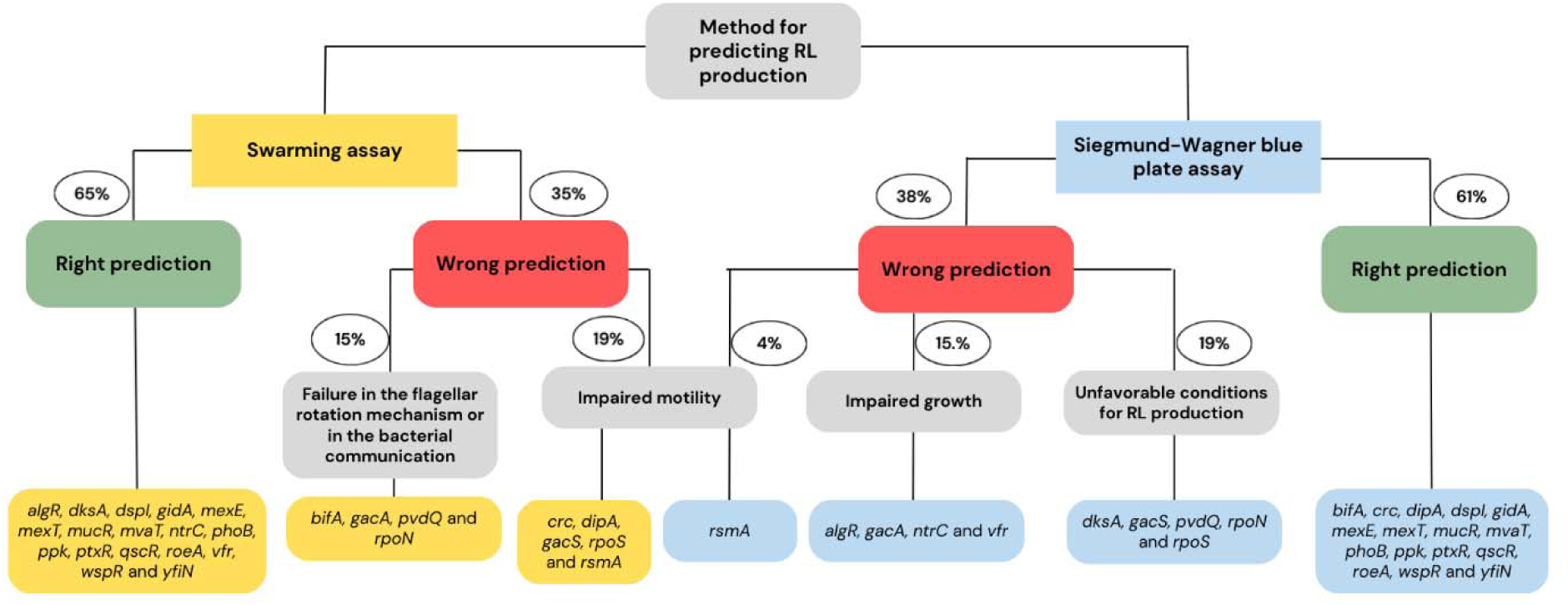
Accuracy of swarming vs. Siegmund-Wagner assays in predicting rhamnolipid production and underlying causes of misprediction.

The swarming assay yielded a 65% rate of correct prediction, confirming that there is some correlation between swarming motility and RL production. The correct predictions were associated with mutations in genes such as *algR, dksA, dspI, gidA, mexE, mexT, mucR, mvaT, rhlC, phoB, ppk, ptxR, qscR, roeA, vfr, wspR,* and *yfiN*. However, 35% of predictions were incorrect, which were attributed to three main biological factors: [1] Failure in the flagellar rotation mechanism or bacterial communication (15%), linked to mutations in *bifA, gacA, pvdQ,* and *rpoN;* [2] Impaired motility, associated with disruptions in *crc, dipA, gacS, rpoS,* and *rsmA;* [3] Loss of motility in the case of the *rsmA* mutation.

The Siegmund–Wagner blue plate assay demonstrated 62% accuracy in predicting RL production, with correct predictions involving the same set of functional genes identified above. Nonetheless, 38% of cases were misclassified, due to: [1] Impaired growth, observed in mutants of *algR, gacA, ntrC,* and *vfr*; [2] Unfavorable conditions for RL production in the assay, involving *dksA, gacS, pvdQ, rpoN,* and *rpoS; and* [3] Impaired motility in *rsmA* mutant.

### Conclusion

Our results reveal that swarming motility and SW blue plates phenotypic assays do not offer a sufficiently reliable prediction of RL production. A considerable proportion of incorrect predictions can occur when relying solely on these methods. While this is not unexpected about swarming, a phenotype also relying on flagellar function, we were surprised by such low reliability for the colorimetric SW blue plates assay. Use of this method should always consider the actual growth and size of the colony formed on the agar surface as influencing the analysis. The observed inconsistencies emphasize the complexity of RL biosynthesis regulation, which includes factors such as QS, stress response, and cellular metabolism. Therefore, while these assays can serve as preliminary screening tools, they must be complemented by direct, quantitative methods, such as LC-MS, to ensure accurate and robust assessment of RL levels.

## Materials and Methods

### Strains

The wildtype strain *P. aeruginosa* PA14 and twenty-nine isogenic knockout mutants in genes previously found or proposed to influence RL production were investigated and are listed in **Table 2**.

**Table 2.**
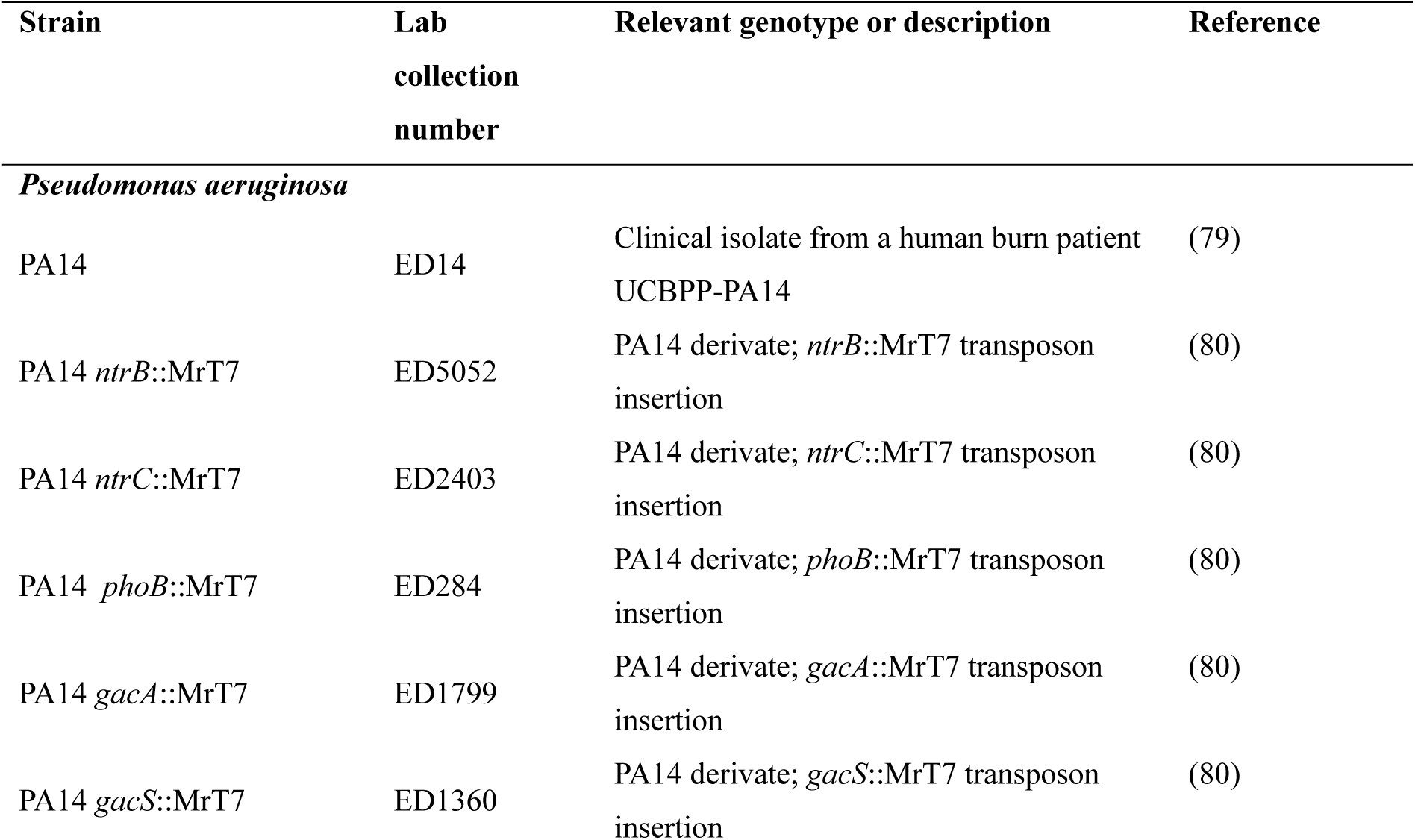
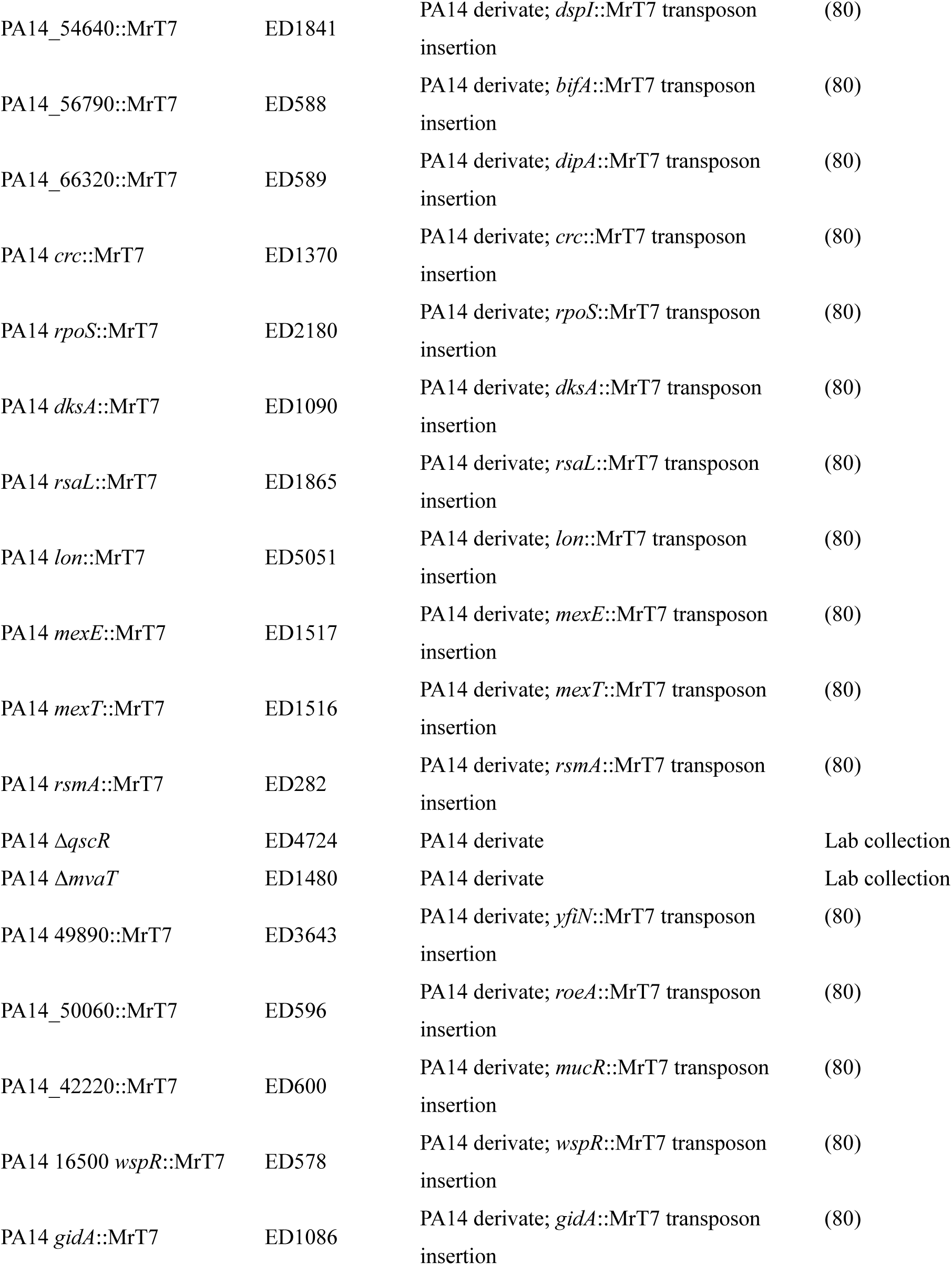
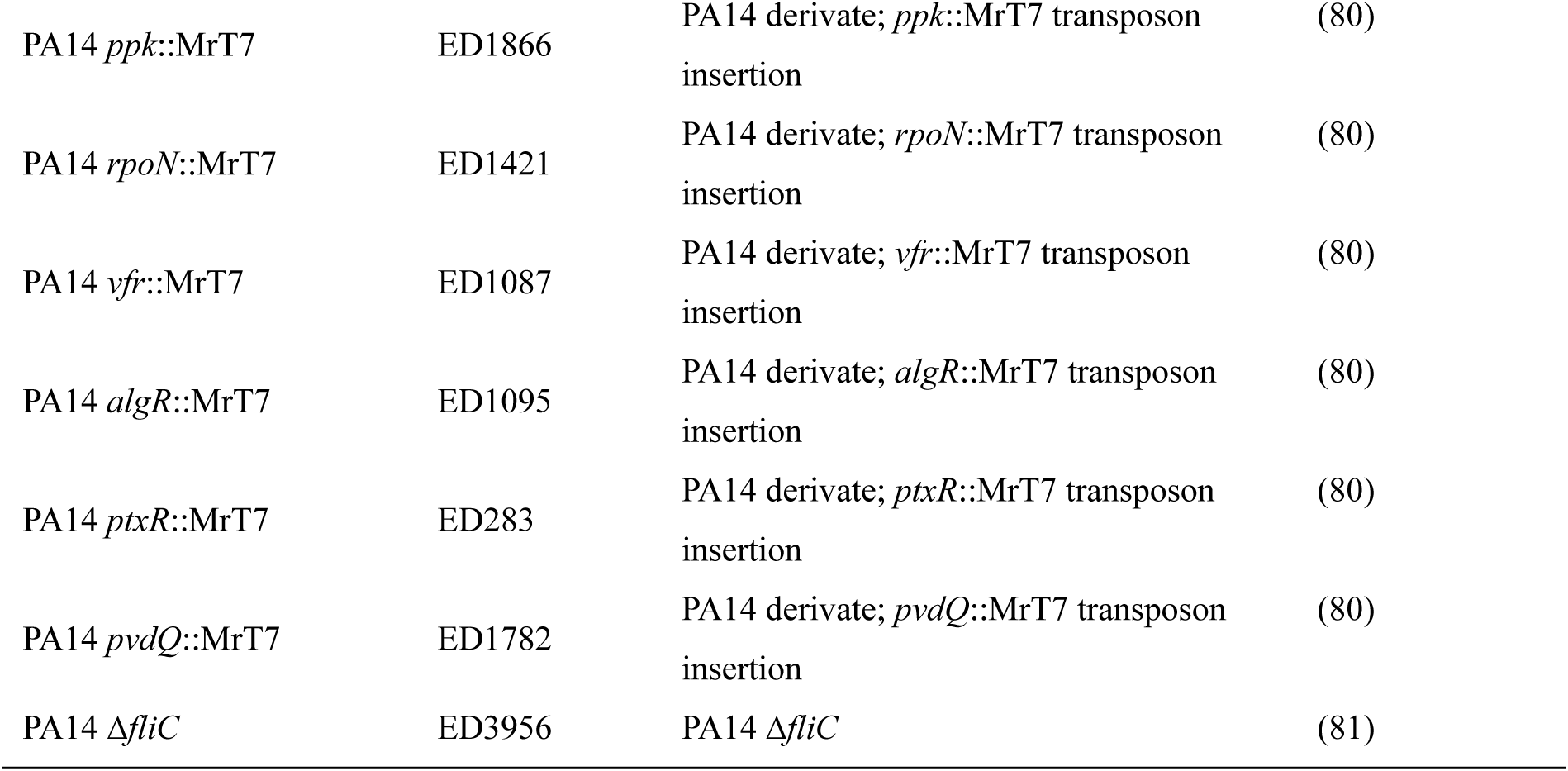
Strains used in this study.

| Strain | Lab collection number | Relevant genotype or description | Reference |
| --- | --- | --- | --- |
| *Pseudomonas aeruginosa* |  |  |  |
| PA14 | ED14 | Clinical isolate from a human burn patient<br>UCBPP-PA14 | (79) |
| PA14 *ntrB*::MrT7 | ED5052 | PA14 derivative; *ntrB*::MrT7 transposon insertion | (80) |
| PA14 *ntrC*::MrT7 | ED2403 | PA14 derivative; *ntrC*::MrT7 transposon insertion | (80) |
| PA14 *phoB*::MrT7 | ED284 | PA14 derivative; *phoB*::MrT7 transposon insertion | (80) |
| PA14 *gacA*::MrT7 | ED1799 | PA14 derivative; *gacA*::MrT7 transposon insertion | (80) |
| PA14 *gacS*::MrT7 | ED1360 | PA14 derivative; *gacS*::MrT7 transposon insertion | (80) |
| PA14_54640::MrT7 | ED1841 | PA14 derivate; *dspI*::MrT7 transposon insertion | (80) |
| PA14_56790::MrT7 | ED588 | PA14 derivate; *bifA*::MrT7 transposon insertion | (80) |
| PA14_66320::MrT7 | ED589 | PA14 derivate; *dipA*::MrT7 transposon insertion | (80) |
| PA14 *crc*::MrT7 | ED1370 | PA14 derivate; *crc*::MrT7 transposon insertion | (80) |
| PA14 *rpoS*::MrT7 | ED2180 | PA14 derivate; *rpoS*::MrT7 transposon insertion | (80) |
| PA14 *dksA*::MrT7 | ED1090 | PA14 derivate; *dksA*::MrT7 transposon insertion | (80) |
| PA14 *rsaL*::MrT7 | ED1865 | PA14 derivate; *rsaL*::MrT7 transposon insertion | (80) |
| PA14 *lon*::MrT7 | ED5051 | PA14 derivate; *lon*::MrT7 transposon insertion | (80) |
| PA14 *mexE*::MrT7 | ED1517 | PA14 derivate; *mexE*::MrT7 transposon insertion | (80) |
| PA14 *mexT*::MrT7 | ED1516 | PA14 derivate; *mexT*::MrT7 transposon insertion | (80) |
| PA14 *rsmA*::MrT7 | ED282 | PA14 derivate; *rsmA*::MrT7 transposon insertion | (80) |
| PA14 $\Delta qscR$ | ED4724 | PA14 derivate | Lab collection |
| PA14 $\Delta mvaT$ | ED1480 | PA14 derivate | Lab collection |
| PA14 49890::MrT7 | ED3643 | PA14 derivate; *yfiN*::MrT7 transposon insertion | (80) |
| PA14_50060::MrT7 | ED596 | PA14 derivate; *roeA*::MrT7 transposon insertion | (80) |
| PA14_42220::MrT7 | ED600 | PA14 derivate; *mucR*::MrT7 transposon insertion | (80) |
| PA14 16500 *wspR*::MrT7 | ED578 | PA14 derivate; *wspR*::MrT7 transposon insertion | (80) |
| PA14 *gidA*::MrT7 | ED1086 | PA14 derivate; *gidA*::MrT7 transposon insertion | (80) |
| PA14 *ppk*::MrT7 | ED1866 | PA14 derivate; *ppk*::MrT7 transposon insertion | (80) |
| PA14 *rpoN*::MrT7 | ED1421 | PA14 derivate; *rpoN*::MrT7 transposon insertion | (80) |
| PA14 *vfr*::MrT7 | ED1087 | PA14 derivate; *vfr*::MrT7 transposon insertion | (80) |
| PA14 *algR*::MrT7 | ED1095 | PA14 derivate; *algR*::MrT7 transposon insertion | (80) |
| PA14 *ptxR*::MrT7 | ED283 | PA14 derivate; *ptxR*::MrT7 transposon insertion | (80) |
| PA14 *pvdQ*::MrT7 | ED1782 | PA14 derivate; *pvdQ*::MrT7 transposon insertion | (80) |
| PA14 Δ*fliC* | ED3956 | PA14 Δ*fliC* | (81) |

### Swarming medium

Swarming motility assays were performed using modified M9 medium with dextrose and casamino acids (M9DCAA) (g.L^-1^): NH_4_Cl – 1.068; K_2_HPO_4_ – 2.99; Na_2_HPO_4_•7H_2_O – 1.70; NaCl – 0.50; Casaminoacids – 5.0; MgSO_4_ 1M - 1000 µL; CaCl_2_ 1M - 1000 µL; dextrose 1.1 M – 10.0 mL and solidified with agar 5 g.L^-1^ Bacto-agar (Difco). The plates containing 20 mL of medium were dried under laminar flow for 75 min, then inoculated with 5.0 µL of suspension prepared in phosphate-buffered saline (PBS) (OD_600_ = 3.0) of the appropriate bacterium and incubated at 37°C (82).

### Bioscreen test

Cells were grown overnight in Tryptic Soy broth (TSB, Difco) at 37°C. Cultures were then inoculated in M9DCAA at an OD_600_ of 0.05 in a final volume of 200 µLin a Honeycomb^TM^ plate. The plate was incubated in a Bioscreen C reader (Growth Curves USA) at 37°C for 36 h with OD_600_ measurements taken every hour, with 10 seconds of shaking prior to each reading.

### Siegmund-Wagner blue agar plates

Medium composition (g.L^-1^): Na_2_HPO_4_ – 0.9; KH_2_PO_4_ - 0.7; CaCl_2_•2H_2_O – 0.1; MgSO_4_•7H_2_O – 0.4; NaNO_3_ – 2.0; tryptone peptone – 1.0; glycerol – 20.0; cetyltrimethyl ammonium bromide (CTAB) – 0.2; methylene blue – 0.01; agar – 20 and trace elements (2.0 mL/L). The trace elements solution consists of (g.L^-1^): FeSO_4•_7H_2_O – 2.0; MnSO4•H_2_O - 1.5; (NH_4_)6Mo_7_O_24•_4H_2_O - 0.6 (69). The pH was adjusted to 7.0 with NaOH solution. The plates were inoculated with 3.0 µL PBS suspensions OD_600_ = 3.0. The plates were incubated at 37°C for 24 h and then maintained at 30°C for 72 h before measurement of the halo zone.

### Swimming medium

Medium composition (g.L^-1^): Tryptic soy broth (TSB) – 25; agar – 2.5. The plates containing 20 mL of medium were dried under laminar flow for 10 min, then inoculated with PBS suspensions OD_600_ = 3.0 using a pipet tip. The plates were incubated at 37°C for 16 h. Colony diameters were measured to calculate the motility area.

### Cultivation conditions and LC-MS quantification of rhamnolipids

Five mL of M9DCAA medium were inoculated at an initial OD_600_ = 0.05 from overnight TSB cultures. Cultures were incubated at 37°C for 120 h in a TC-7 roller drum (New Brunswick Scientific) at 240 rpm. At the end of the incubation period, 1.0 mL of culture was collected and centrifuged at 10,000 rpm for 5m. The supernatant was used for RL quantification using LC-MS, whereas the pellet was dried and weighed to determine biomass dry weight. For LC-MS analysis, the supernatant was diluted in acetonitrile supplemented with 7.9 ppm 16-hydroxyhexadecanoic acid as an internal standard. LC-MS measurements were performed on a Waters Quattro Premier XE triple quadrupole mass spectrometer coupled to a Waters 2795 HPLC system (Waters, Mississauga, ON). Chromatographic separation was achieved on a Phenomenex Kinetex C8 column (2.6 µm, 100 × 4.6 mm) under isocratic elution with water and acetonitrile containing 1% formic acid, as previously described (83).

### Statistical Analysis

Statistical analyses were performed using GraphPad Prism. Data was analyzed by ordinary one-way ANOVA followed by Dunnett’s post hoc test for multiple comparisons.

## Acknowledgments

The authors are grateful to Marie-Christine Grouleau for critically reading the manuscript and providing helpful comments. This research was financially supported by FAPESP (São Paulo Research Foundation) operating grant 2024/01150-0 and by Discovery Grant RGPIN-2020-06771 from the Natural Sciences and Engineering Research Council of Canada (NSERC).

